# Elysium: RNA-seq Alignment in the Cloud

**DOI:** 10.1101/382937

**Authors:** Alexander Lachmann, Zhuorui Xie, Avi Ma’ayan

**Affiliations:** Department of Pharmacological Sciences, Icahn School of Medicine at Mount Sinai, One Gustave L. Levy Place, Box 1603, New York, NY 10029, USA; Big Data to Knowledge, Library of Integrated Network-based Cellular Signatures, Data Coordination and Integration Center (BD2K-LINCS DCIC); Mount Sinai Knowledge Management Center for Illuminating the Druggable Genome (KMC-IDG); Team Nitrogen, NIH Data Commons Pilot Project Consortium (DCPPC)

## Abstract

**Motivation:** RNA-sequencing (RNA-seq) is currently the leading technology for genome-wide transcript quantification. Mapping the raw reads to transcript and gene level counts can be achieved by a variety of aligners and pipelines. The diversity of processing options reduces interoperability. In addition, the alignment step requires significant computational resources and basic programming knowledge. Elysium enables users of all skill levels to perform a uniform and free RNA-seq alignment in the cloud.

**Results:** The Elysium infrastructure is comprised of four components: A file upload API that enables storage of FASTQ files on Amazon S3 without Amazon credentials; an API to handle the cloud alignment job scheduling for uploaded files; and a graphical user interface (GUI) to provide intuitive access to users that do not have command-line access skills.

**Availability:** The Elysium source code is available under the Apache Licence 2.0 on GitHub at: https://github.com/maayanlab/elysium

The service of cloud based RNA-seq alignment is freely accessible through the Elysium GUI at: http://elysium.cloud

## Background

Genome-wide gene expression data from thousands of studies have been accumulating and made available for exploration and reuse through public repositories such as the Gene Expression Omnibus (GEO) (1). RNA-seq is supplanting cDNA microarrays as the dominant technology due to competitive cost, increased sensitivity, ability to quantify splice variants, and improved reproducibility. The processing of RNA-seq data remains a challenge due to computational resource demand as well as the required technical skills needed to execute computational pipelines. The process of aligning raw reads to a reference genome can be performed with a multitude of alignment algorithms (2–6) as well as different genome assemblies and transcript annotations. Several attempts to reprocess large collections of publicly available samples address the need for uniform processing for better interoperability (7, 8). In this context, we developed ARCHS4 (9). ARCHS4 processes most of the RNA-seq data from GEO collected from human or mouse, supplying the largest repository of homogeneously processed RNA-seq data with 149,222 mouse and 125,501 human samples currently available. ARCHS4 has a scalable cloud computing infrastructure capable to adjust to fluctuations in demand. Efficient implementation of storage and cloud compute instances are critical for a cost efficient and scalable solution. Here we present a new API driven implementation that orchestrates four components to allow users to align their own RNA-seq data in the cloud with an enhanced version of the ARCHS4 pipeline.

**Fig.1.**
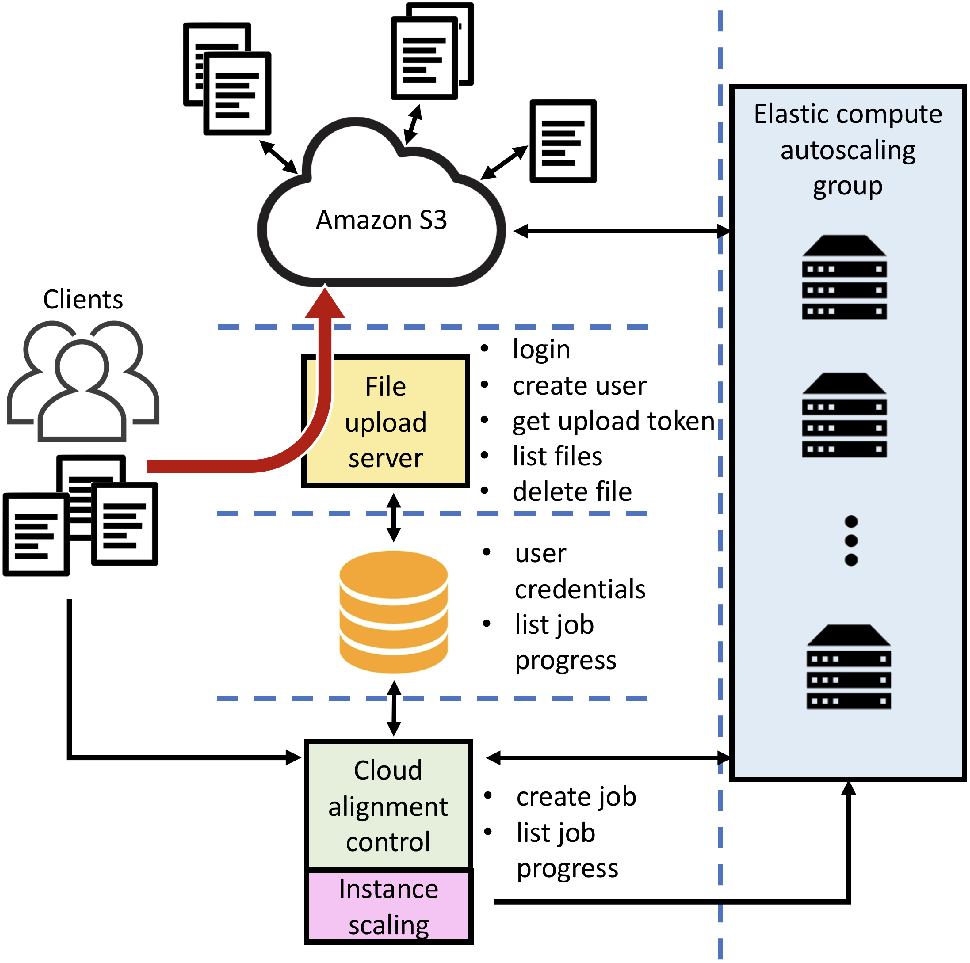
The Elysium API RNA-seq architecture. Users upload data into Amazon S3 using the file upload server API. Once FASTQ files are deposited in a cloud storage, alignment jobs can be scheduled through the cloud alignment control server API. The elastic back-end initiates Docker containers on request and deposits gene counts and transcript counts to the S3 user repository.

## Materials and methods

The Elysium APIs are cloud-based and built for scalability. To achieve this, critical aspects of the pipeline are averting bottlenecks, for example, relying on individual servers for handling heavy duty tasks such as file upload and data processing. The system relies on control servers that upon request produce temporary user credentials for allowing direct interaction with the Amazon S3 bucket for file upload and submission of job instructions to the compute backend. The system is composed of the following four components:

### Upload

The file upload server is hosting an API to facilitate file upload directly to the Amazon S3 bucket without passing data through an intermediary instance. In order to upload a file, a user needs to register for a user account. With this account, encrypted tokes can be requested. Using tokens, POST requests are sent directly to Amazon S3 without the need for Amazon credentials. The RNA-seq data is stored in a personal repository.

### Job

The job scheduler API supports submission of alignment jobs for files already uploaded to the cloud repository. The user can specify a single, or paired end, read files, and the reference genome for the alignment. The submitted jobs are moved into a processing queue that serves the job based on its description on a first-come-first-serve basis. The job scheduler also handles instance scaling of the processing backend. It can be configured with a minimum and a maximum number of active compute instances, as well as a scaling factor relative to the number of waiting alignment jobs.

### AWSalign

The alignment container is a preconfigured Docker (10) container that is automatically deployed into the elastic Amazon cluster. The container runs Kallisto (11) as the main alignment tool. The alignment tool performs transcript quantification and gene count transformation, and then moves the alignment results to the user workspace. After a job is completed, the alignment container will request a new job from the job scheduler API.

### GUI

The Elysium GUI is a light-weight graphical userinterface that is serving the underlying API functionality. It supports the generation of user accounts, file upload, job generation and submission. Elysium displays job processes and results in the user workspace with download links for transcript level quantification and gene level count processed data files.

## Implementation

The file upload and job scheduler APIs are dockerized tornado web-servers. The Docker containers are fully configurable through passing environment parameters during the Docker container launch. User credentials and job descriptions are stored in a MySQL database shared between the file upload server and the job scheduler. The AWSalignment container is a dockerkized Ubuntu server supporting python and R with a MySQL client. AWSalignment is preconfigured with Kallisto 0.44 for sequence alignment. For the transfer of transcript quantification results back to the user’s personal account on Amazon S3, the python boto library is used. Transcript to gene mapping is performed with an R-script, and the annotation from the cDNA annotation file from Ensembl. The annotation version is Ensembl 90, and the reference genome for human and mouse are GRCh38 and GRCm38, respectively. These files are downloaded from the Ensembl website. The Elysium GUI is implemented as a single HTML file. The GUI is an interactive user interface. This is achieved with support from the JQuery, Bootstrap, and CSSgrid libraries. The page is hosted via the job scheduler tornado server. APIs are registered and documented with SwaggerHub and SmartAPI.

## Conclusion

Elysium enables fast and scalable RNA-seq alignment in the cloud. Elysium has native programmatic access through the API to its functionality and alternative GUI. Through automatic instance scaling of the compute back-end, the maintenance and operating costs of the cloud alignment jobs is kept low. The average cost for processing a FASTQ file is currently below ¢1 as previously reported (9). A key capability of Elysium is the leveraging of many of the available features of the cloud infrastructure, and as a consequence minimizing possible bottlenecks in data transfer and computational load. Upload speed per file is currently at 60MB/s and this speed is independent of concurrent users. The dynamic cluster size and fast load times of Docker containers result in a scaling response time of fully functional resources in less than 5 minutes. This keeps wait times from submission to final results on average below 15 minutes, and this waiting time is independent of traffic. In summary, Elysium is an example of an application that leverages cloud computing to provide API services. Applications other than RNA-seq analysis could be incorporated into the existing Elysium framework with the addition of other types of specialized Docker containers. The APIs presented here, in combination with the GUI, increase interoperability of existing RNA-seq data analysis pipelines, facilitating users with minimal computational skills with the ability to process their own data using the same pipeline implemented for creating the ARCHS4 resource. The uniform processing can place the newly processed data in context of more than 250,000 previously published RNA-seq data-sets currently available at the ARCHS4 resource.

## ACKNOWLEDGEMENTS

This work is partially supported by the National Institutes of Health (NIH) grants U54-HL127624 (LINCS-DCIC), U24-CA224260 (IDG-KMC), and OT3-OD025467 (NIH Data Commons), as well as cloud credits from the NIH BD2K Commons Cloud Credit Pilot project to AM.

## Bibliography

1. Ron Edgar, Michael Domrachev, and Alex E Lash. Gene expression omnibus: Ncbi gene expression and hybridization array data repository. Nucleic acids research, 30(1):207–210, 2002.

2. Ben Langmead and Steven L Salzberg. Fast gapped-read alignment with bowtie 2. Nature methods, 9(4):357–359, 2012.

3. Ruiqiang Li, Chang Yu, Yingrui Li, Tak-Wah Lam, Siu-Ming Yiu, Karsten Kristiansen, and Jun Wang. Soap2: an improved ultrafast tool for short read alignment. Bioinformatics, 25 (15):1966–1967, 2009.

4. Alexander Dobin, Carrie A Davis, Felix Schlesinger, Jorg Drenkow, Chris Zaleski, Sonali Jha, Philippe Batut, Mark Chaisson, and Thomas R Gingeras. Star: ultrafast universal rna-seq aligner. Bioinformatics, 29(1):15–21, 2013.

5. Heng Li and Richard Durbin. Fast and accurate short read alignment with burrows–wheeler transform. Bioinformatics, 25(14):1754–1760, 2009.

6. Chi-Man Liu, Thomas Wong, Edward Wu, Ruibang Luo, Siu-Ming Yiu, Yingrui Li, Bingqiang Wang, Chang Yu, Xiaowen Chu, Kaiyong Zhao, et al. Soap3: ultra-fast gpu-based parallel alignment tool for short reads. Bioinformatics, 28(6):878–879, 2012.

7. Leonardo Collado-Torres, Abhinav Nellore, Kai Kammers, Shannon E Ellis, Margaret A Taub, Kasper D Hansen, Andrew E Jaffe, Ben Langmead, and Jeffrey T Leek. Reproducible rna-seq analysis using recount2. Nature Biotechnology, 35(4):319–321, 2017.

8. John Vivian, Arjun Arkal Rao, Frank Austin Nothaft, Christopher Ketchum, Joel Armstrong, Adam Novak, Jacob Pfeil, Jake Narkizian, Alden D Deran, Audrey Musselman-Brown, et al. Toil enables reproducible, open source, big biomedical data analyses. Nature biotechnology, 35(4):314, 2017.

9. Alexander Lachmann, Denis Torre, Alexandra B Keenan, Kathleen M Jagodnik, Hoyjin J Lee, Lily Wang, Moshe C Silverstein, and Avi Ma’ayan. Massive mining of publicly available rna-seq data from human and mouse. Nature communications, 9(1):1366, 2018.

10. Dirk Merkel. Docker: Lightweight linux containers for consistent development and deployment. Linux J., 2014(239), March 2014. ISSN 1075–3583.

11. Nicolas L Bray, Harold Pimentel, Páll Melsted, and Lior Pachter. Near-optimal probabilistic rna-seq quantification. Nature biotechnology, 34(5):525, 2016.

